# In silico evaluation of the role of the long non-coding RNA LINC00092 in thyroid cancer progression ; regulation of the miR-34a-5p/RCAN1 axis

**DOI:** 10.1101/2022.09.04.506551

**Authors:** Saman Morovat, Pejman Morovat, Mohammad Javad Kamali, Shahram Teimourian

**Author notes:** **Corresponding author: Shahram Teimourian**, Ph.D.; Department of Medical Genetics and Molecular Biology, Faculty of Medicine, Iran University of Medical Sciences (IUMS), Tehran, Iran.

## Abstract

**Background:** As the most prevalent endocrine cancer, thyroid cancer (TC) accounts for 1.7% of all cancer cases. A significant increase in TC morbidity has been observed over the past three decades. TC diagnosis has been reported to be problematic based on the current approach. As a result, it is imperative to develop molecular biomarkers to improve the accuracy of the diagnosis. An analysis of bioinformatics data was conducted in this study to analyze lncRNAs and their roles as ceRNAs associated with the development and progression of TC.

**Materials and Method:** The first step in this study was to collect RNA-seq data from the GDC database. Then, DESeq2 was used to analyze differentially expressed lncRNAs (DElncRNAs), miRNAs (DEMIs), and mRNAs (DEGs) between TC patients and healthy subjects. Our study identified DElnc-related miRNAs and miRNA-related genes to develop a lncRNA/miRNA/mRNA axis using online tools and screening. A co-expression analysis was performed to investigate correlations between DElncs and their associated mRNAs. Next, a protein-protein interaction (PPI) network was constructed. Functional enrichment and pathway enrichment were conducted on genes in the PPI network to discover additional biological activities among these molecules. Lastly, a correlation between the expression levels and the infiltration abundance of immune cells was assessed through immune infiltration analysis.

**Results:** There were 58 DElncs, 34 DEMIs, and 864 DEGs in thyroid tumor tissue and non-tumor tissue samples. Following validation of our lncRNA results with the intersection of differentially expressed lncRNAs in TCGA and GEPIA2, we selected two downregulated DElncs, including AC007743.1 and LINC00092, as the final research elements. We then performed an interaction analysis to predict lncRNAs-miRNAs and miRNAs-mRNAs interactions, which led to identifying the LINC00092/miR-34a-5p and miR-34a-5p/RCAN1 axis, respectively. There was a correlation between LINC00092 and RCAN1 according to Pearson correlation analysis. To improve our understanding of RCAN1, we developed a PPI network. According to the Immune Infiltration Analysis, RCAN1 expression was positively correlated with CD8+ T cells, macrophages, and neutrophils.

**Conclusion:** The results of this study suggest that LINC00092/miR-34a-5p/RCAN1 axis may have a functional role in the progression of TC. LINC00092 may be used as a promising biomarker for TC prognosis and may be a better diagnostic and therapeutic target.

## Introduction

Thyroid cancer (TC) is the most common endocrine malignancy and is characterized by an enlarged thyroid gland in the neck (1). This disease accounts for 1.7% of all new cancer cases, and its annual new case detection rate has been increased due to the undisputable developments and breakthroughs in the diagnostic techniques (2). The risk of developing this form of cancer is three times higher in women than in men (3). Due to the steady increase in its incidence, it is predicted that by 2030 it will overtake breast cancer as the third most prevalent type of cancer among women (4,5). Over the past three decades, there has been a significant increase in TC morbidity (6). It has been estimated that 10% of TC patients have malignant tumors (7), and approximately 7% of TC patients lost their lives in 2020 (8).

Patients’ age is crucial in cancer subtype diagnosis (4). Ultrasound examination is the preferred diagnosis method, while percutaneous fine-needle aspiration biopsy (PFNA) is the standard gold test for clarifying benign from malignant thyroid nodules (9). PFNA’s notable dependency on the pathologists’ diagnostic expertise and specimen size, significantly limits its functionality. However, advances in molecular medicine have expanded treatment choices (10).

An epigenetic modification results in inherited phenotypic changes to a gene without altering its DNA sequence; these changes may have significant consequences on the development of TCs (11). The four major epigenetic modifications are DNA methylation, chromatin remodeling, histone modification, and non-coding RNA regulation(12).

The human genome consists of 95% non-coding DNA. The transcription of more than 10,000 different non-coding RNAs (ncRNAs) from non-coding DNAs has been found to include microRNAs (miRNAs), small interfering RNAs (siRNAs), asRNAs, and long non-coding RNAs (lncRNAs). Recent studies have linked regulatory interactions between non-coding RNAs and thyroid cancer progression, but their role remains unclear (13). The competing endogenous RNA (ceRNA) hypothesis was introduced by Leonardo Salmena and colleagues in 2011 (14). This hypothesis suggests that ceRNAs are partial transcription products that regulate various biological processes. Among these regulatory molecules are messenger RNAs (mRNAs), circular RNAs (circRNAs), long non-coding RNAs (lncRNAs), and pseudogenic RNAs. It has been shown that these transcripts compete for miRNAs and regulate each other post-transcriptionally. Multiple diseases originated from the disruption of this network, such as coronary artery disease, neurodegenerative diseases, and cancers of various types (15–17). Many studies have shown that thyroid cancer’s development, invasion, and metastasis are associated with the dysregulation of ceRNA networks (18). Developing effective diagnostic procedures and treatment options for thyroid cancer stages requires a deep understanding of these dysregulations.

LncRNAs are non-coding RNAs with at least 200 base pairs (bp) (19), which are thought to regulate gene expression at the transcriptional and post-transcriptional levels, among many other functions (20,21). LncRNAs may have a tumor-suppressive or oncogenic role in human malignancies like TC, which is why they have been proposed as prognostic, diagnostic, and therapeutic biomarkers in cancer research.

Zhou Y et al. showed that MAPKAPK5-AS1 enhanced thyroid cancer cell proliferation and migration by serving as a sponge for miR-519e-5p by regulating YWHAH expression (22). On the other hand, other studies substantiated the role of lncRNAs as tumor suppressors. According to research by Wang et al., MEG3 inhibits tumor invasion and migration by targeting the RAC1 gene (23).

In 2005, National Cancer Institute and the National Human Genome Research Institute launched the Cancer Genome Atlas Project as a three-year pilot project to study lung, brain, and ovarian cancers (24). Thousands of tumor samples representing more than twenty types of cancer have been collected as part of this project over the years. By improving cancer therapies, diagnosis, and prevention measures, this project will pave the way for discoveries and novel methods.

Our study obtained LncRNAs, miRNAs, and mRNA sequencing data for Thyroid Carcinoma (THCA) from the Cancer Genome Atlas database (TCGA). Differential expression analysis was then performed using R packages to detect lncRNAs (DElncs), miRNAs (DEMIs), and mRNAs (DEGs). In addition, online tools and screening were utilized to obtain DElncs-related miRNAs and miRNA target genes to develop a lncRNA/miRNA/mRNA axis. As part of the study, a co-expression analysis was performed to investigate correlations between DElncs and their associated mRNAs. Then, a protein-protein interaction (PPI) network for the final target gene was developed. Functional enrichment and pathway enrichment were conducted on genes in the PPI network to discover additional biological activities among these molecules. For the final step, an immune infiltration analysis was performed to determine whether a correlation existed between the expression levels of our final target and the infiltration abundance of immune cells (Fig 1).

**Fig 1.**
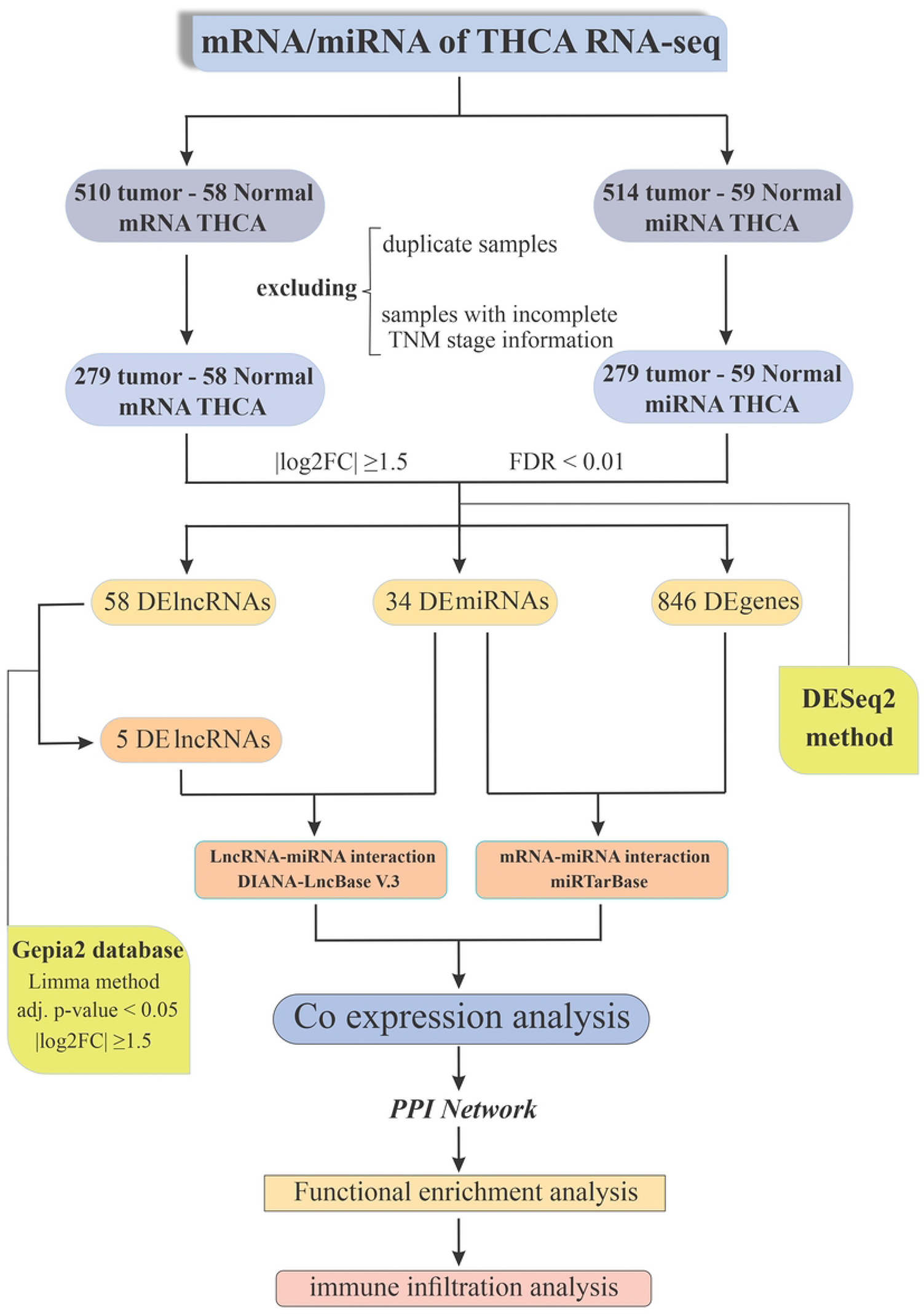
Flowchart depicts the procedures used in this study to develop the final regulatory axis for thyroid cancer.

## Materials and Methods

### RNA Sequencing Data Preprocessing

On March 8, 2022, the GDC data portal (https://portal.gdc.cancer.gov/) related to the TCGA project was used to obtain RNA-Seq, miRNA-Seq expression data, and clinical information for thyroid carcinoma (project ID= TCGA-THCA). There were 510 primary tumors, 58 adjacent non-tumor tissues for lncRNAs and mRNAs, 514 primary tumors, and 59 adjacent non-tumor tissues for miRNAs. After excluding duplicate samples and samples with incomplete clinical-TNM stage information, the final number of RNA samples in the primary tumor and adjacent non-tumor tissue decreased to 279 (203 females and 76 males) and 58 (41 females and 17 male), respectively. Also, the final number of miRNA samples in the primary tumor and adjacent non-tumor tissue decreased to 279 (203 females and 76 males) and 59 (42 females and 17 males), respectively (Table 1).

**Table 1.**
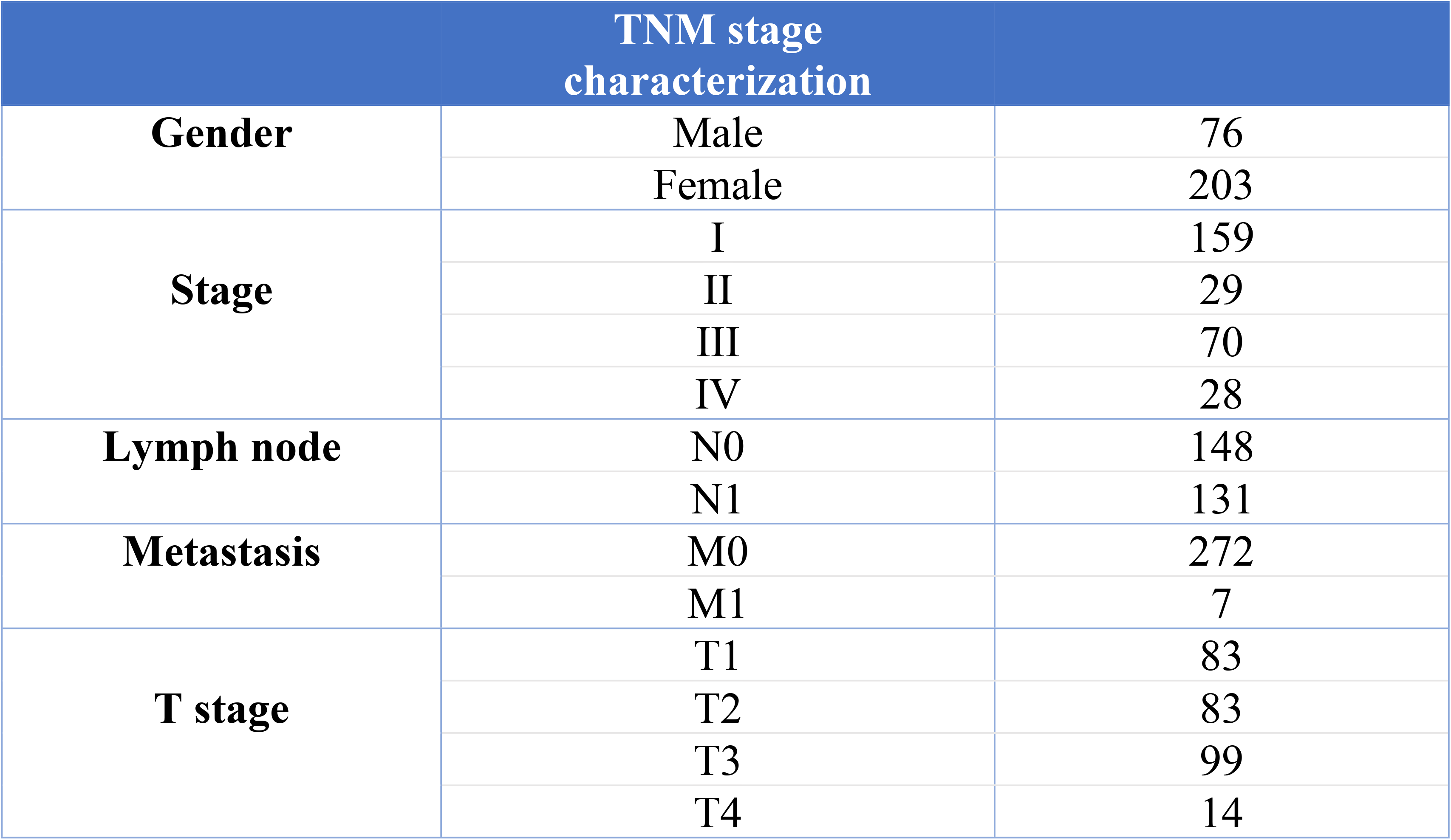
TNM stage characteristics of patients with thyroid cancer from TCGA.

| TNM stage characterization |  |  |
| --- | --- | --- |
| Gender | Male | 76 |
|  | Female | 203 |
| Stage | I | 159 |
|  | II | 29 |
|  | III | 70 |
|  | IV | 28 |
| Lymph node | N0 | 148 |
|  | N1 | 131 |
| Metastasis | M0 | 272 |
|  | M1 | 7 |
| T stage | T1 | 83 |
|  | T2 | 83 |
|  | T3 | 99 |
|  | T4 | 14 |

### Differential expression lncRNAs (DElncs) of THCA data and Gepia2

The first step in this process was the standardization by the normalization of data. Normalization was performed with the GDCRNAtools package (25) in R software using TMM (26) and Voom (27) methods. We analyzed the lncRNA expression data between the primary tumor and normal solid tissue using the DESeq2 package (V 1.34.0) (28) in the R programming language to identify differentially expressed lncRNAs (DElncs). DElncs were detected based on |log2foldchange| > 1.5 and FDR < 0.01 criteria.

GEPIA2 (29) is an improved version of GEPIA that uses a standard processing pipeline to analyze the RNA sequencing expression data of 9,736 tumors and 8,587 normal samples from the TCGA and GTEx studies. Several customizable functions are available in GEPIA2, such as tumor/normal differential expression analysis, cancer type or pathological stage profiling, patient survival analysis, similar gene discovery, correlation analysis, and dimensionality reduction analysis. The GEPIA2 database was used in this study to further analyze differentially expressed lncRNAs in THCA and validate our results. The expression of these genes was analyzed using the limma method (30), with the following filter criteria: |log2foldchange| >1.5 and FDR < 0.01. Our study performed an intersection between the results of differentially expressed lncRNAs in the TCGA database and those in the GEPIA2 dataset.

### Differential expression miRNAs (DEMIs) and mRNAs (DEGs) of THCA data

The DESeq2 package was also used to screen differentially expressed miRNAs (DEMIs) and differentially expressed mRNAs (DEGs) in the TCGA dataset between normal and tumor samples. DESeq2 computes the geometric mean for each gene across all samples as an internal normalization method. After doing this, the number of genes in each sample was divided by the mean. A threshold of |log2FoldChange| >1.5, and FDR < 0.01 were set in this case to detect DEMIs and DEGs. We considered this criterion to be statistically significant.

### Prediction of lncRNA-miRNA interactions (MREs)

DIANA-LncBase version 3 (http://diana.e-ce.uth.gr/lncbasev3) repository (31) was used to predict miRNA response element (MRE) sites for each DElnc (lncRNA–miRNA interaction). DIANA-LncBase v3 is a database providing miRNA targets on non-coding transcripts that have been experimentally validated. Approximately 500,000 entries in the DIANA-LncBase v3 database correspond to ∼240,000 distinct tissue and cell type-specific miRNA-lncRNA interactions. The interactions are specified by 15 low- and high-throughput techniques, corresponding to 243 different cell types/tissues and 162 different experimental conditions.

Then, to identify potential target miRNAs on DElncs, an intersection was performed between DIANA-LncBase targeted miRNAs and dysregulated DEMIs from TCGA results.

### Prediction of miRNA-mRNA interactions

The package multiMiR (32) in the R programming language was used to obtain miRNA target genes (miRNA-mRNA interactions). MultiMiR with a web server at (http://multimir.org/) provides a comprehensive collection of predicted and validated miRNA-target interactions as well as their correlations with diseases and drugs. Among its many features, multiMiR has a compilation of 14 separate databases, the most complete than any other collection. Also, an expansion of databases to those based on disease annotation and drug response, in addition to many experimental and computational databases, by user-defined cutoffs for predicted binding strength, provides the most confident selection.

This study used miRTarBase (33), a validated database from multiMiR. Next, with scripts written in R, predicted mRNAs of downregulated miRNAs acquired from miRTarBase were merged with upregulated DEGs from TCGA results. In contrast, predicted mRNAs of upregulated miRNAs were merged with downregulated DEGs.

### Co-expression analysis

The correlation between DElncs and their related mRNAs was determined using the GDCRNATools package in the R programming language. The minimum interaction absolute value was set at medium confidence 0.300 with a P-value cutoff <0.001. Based on the findings, Protein-protein interaction analysis was performed on the correlated genes for further validation.

### Protein-protein interaction (PPI) analysis

A PPI (Protein-Protein Interaction) network associated with DElncs-targeted mRNAs was constructed using GeneMANIA (34), a plugin in Cytoscape software (35). A GeneMANIA platform can be used to predict protein-protein interactions, protein-DNA, and genetic interactions, pathways analysis, protein expression inspection, protein domains, and phenotypic screening profiles. Three prominent cases are provided in GeneMANIA: genetic queries, genetic network searches, and gene-by-gene queries.

According to Gene Ontology (GO) network weighting, the top 20 related genes were identified based on co-expression, co-localization, genetic interactions, pathways, physical interactions, and predicted and shared protein domains.

### Functional Enrichment Analysis

Using the clusterProfiler package (36) in R, we investigated the function and pathway information enriched for DElncs-targeted mRNAs based on gene ontology (GO) (37) and Kyoto Encyclopedia of Genes and Genomes (KEGG) (38) information. Clusterprofiler identified enriched genes in PPI networks based on hypergeometric distribution tests using groupGo, enrichGO, and enrichKEGG. Functional annotation with p-value < 0.05 and q-value < 0.05 was considered statistically significant.

### Immune Infiltration Analysis

We also investigated the correlation between our final target gene expression levels and the infiltration abundance of six immune cells (B cells, CD4 + T cells, CD8+ T cells, neutrophils, macrophages, and dendritic cells), which were estimated by TIMER (version 2.0; https://cistrome.shinyapps.io/timer/) (39). The TIMER algorithms assessed the correlation between final target gene expression and immune infiltration in the THCA tumors. P-values < 0.05 and −1< R < 1 were obtained with the Spearman’s rank correlation test and were considered statistically significant.

## Results

### Identification of differential expressed lncRNAs

The DESeq2 package in R was used to identify differentially expressed lncRNAs (DElncs) in the thyroid cancer and normal control samples. DElncs comprised 48 upregulated genes and 10 downregulated genes (Fig 2A). When we merged DElncRNAs from the TCGA and GEPIA2 databases results, only five DElncs remained, including AC007743.1, DOCK9-AS2, LINC00092, MIR31HG, and TNRC6C-AS1. After further analysis and filtration of lncRNAs that have already been studied in thyroid cancer, among these five DElncs, two downregulated DElncs, including AC007743.1 and LINC00092, were chosen as the final research elements in this study (Table 2).

**Table 2.** The expression analysis results for five long noncoding RNAs were obtained by merging the TCGA and Gepia2 results.

| TCGA Results |  |  |  | Gepia2 Results |  |  |
| --- | --- | --- | --- | --- | --- | --- |
| Symbol | log2FC | p-value | FDR | Gene.ID | Log2FC | Adj. p-value |
| AC007743.1 | -1.55 | 4.15E-31 | 5.58E-30 | ENSG00000233251.7 | -1.94 | 1.5E-211 |
| DOCK9-AS2 | 1.96 | 6.43E-61 | 3.40E-59 | ENSG00000260992.1 | 1.56 | 1E-102 |
| LINC00092 | -1.65 | 2.07E-20 | 1.44E-19 | ENSG00000225194.2 | -1.73 | 1.1E-124 |
| MIR31HG | 2.91 | 3.40E-42 | 8.21E-41 | ENSG00000171889.3 | 1.56 | 1.84E-76 |
| TNRC6C-AS1 | 2.58 | 9.59E-67 | 6.11E-65 | ENSG00000204282.4 | 1.67 | 5.64E-61 |

**Fig 2.**
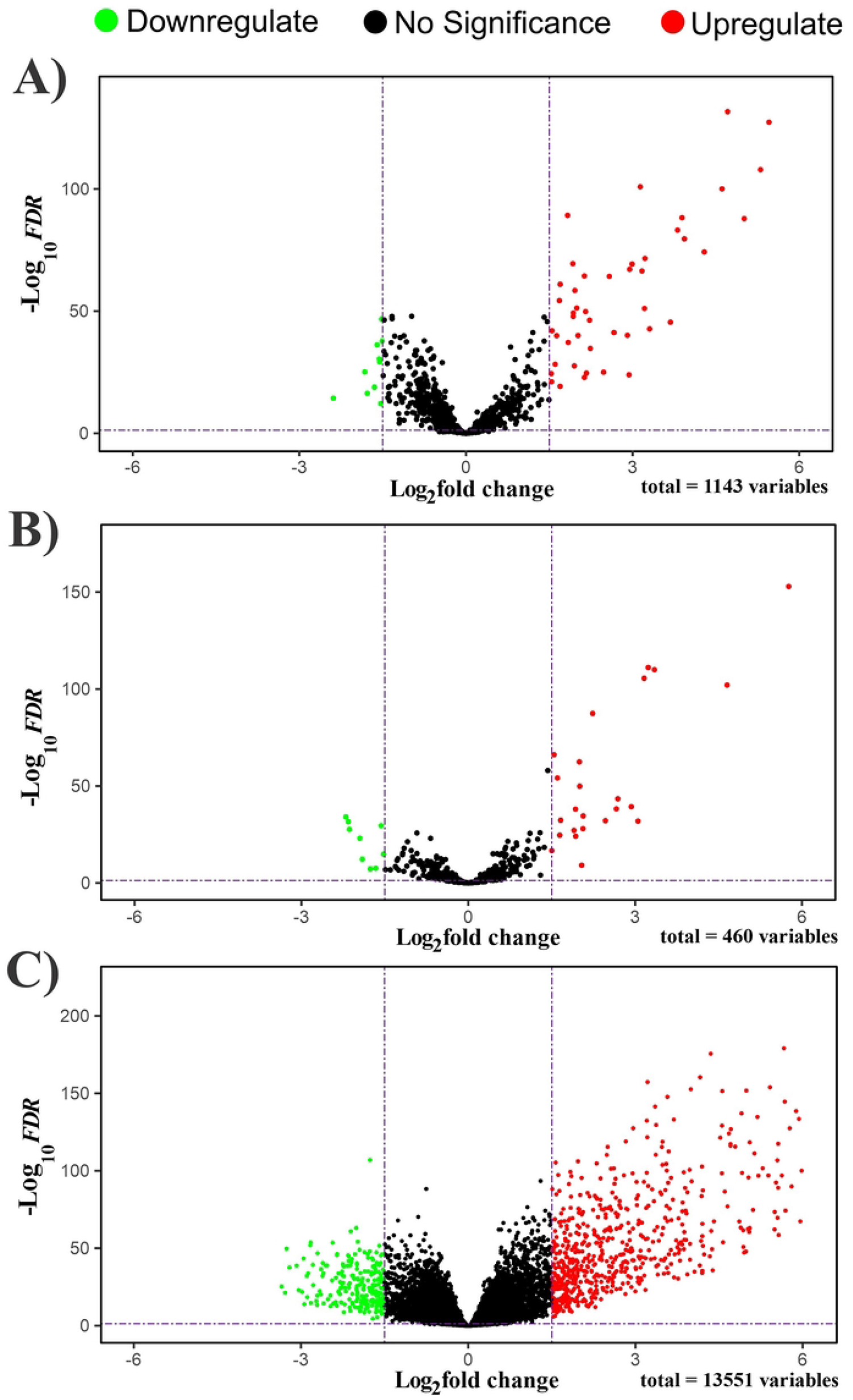
Volcano plots of differentially expressed **A)** lncRNAs (DElncs), **B)** miRNAs (DEMIs), and **C)** mRNAs (DEGs). In tumor samples, green dots represent DE RNAs that have been downregulated, and red dots represent DE RNAs that have been upregulated. Black dots indicate RNAs that have been excluded.

### Identification of differentially expressed miRNAs and mRNAs

The DEMIs (Differentially expressed miRNAs) and DEGs (Differentially expressed genes) between primary tumors and normal solid tissue were identified using the DESeq2 package analysis performed on samples taken from the TCGA-THCA database. The Volcano map shows that DEMIs consist of 25 upregulated and 9 downregulated genes, whereas DEGs consist of 638 upregulated genes and 226 downregulated genes. For the determine determination of DEGs and DEMIs, these criteria were used: |log2foldChange| > 1.5 and FDR < 0.01 (Fig 2, B, C).

### Prediction of lncRNA-miRNA interactions (MREs)

It has been increasingly demonstrated that lncRNAs play a vital role in tumors by acting as a “sponge” to absorb miRNAs. In this way, some miRNAs related to these two DElncs (AC007743.1 and LINC00092) were predicted according to the ceRNA hypothesis. In order to investigate whether these two DElncs function as ceRNA in TC, the DIANA-LncBase v3 database was used to collect potential target miRNAs. Considering that the DElncs chosen for this study were downregulated, and lncRNAs lead to miRNA upregulation, we intersected the predicted miRNAs by DIANA-LncBase v3 with 25 upregulated DEMIs from the TCGA-THCA analysis results. In this way, 2 miRNA-lncRNA interactions were discovered, including AC007743.1/hsa-miR-221-3p and LINC00092/hsa-miR-34a-5p.

### Prediction of miRNA-mRNA interactions

MiRNAs can bind to complementary sequences inside mRNAs (MREs), which can lead to their degradation or suppression; therefore, we have used the multiMiR package to predict which RNA has complementary binding sites for our two miRNAs (hsa-miR-221-3p, hsa-miR-34a-5p), based on miRTarBase databases.

CeRNA hypothesis states that downregulating DElncs results in upregulating corresponding target miRNAs, and since more miRNAs are present in the cytoplasm, a greater percentage of target gene transcripts is degraded and suppressed. Therefore, an intersection between targeted mRNAs obtained by multiMiR and downregulated DEGs from TCGA was carried out. By this approach, miR-34a-5p/RCAN1interaction was identified by validated experimental methods in miRTarBase database like Immunohistochemistry, Luciferase reporter assay, qRT-PCR, and Western blot, which are strong evidence.

### Co-expression analysis between lncRNAs and mRNAs

The correlation analysis between LINC00092 and RCAN1 is presented in Figure 3 based on the predicted Pearson correlation score. Significant correlations were found between LINC00092 and RCAN1 (Cor=0.37, P= 1.16e-12). The final axis selected for more analysis and investigation was LINC00092/hsa-miR-34a-5p/RCAN1. All the expression information of our final axis separately (LINC 00092/hsa-miR-34a-5p/RCAN1) have shown in the boxplot (Fig 4).

**Fig 3.**
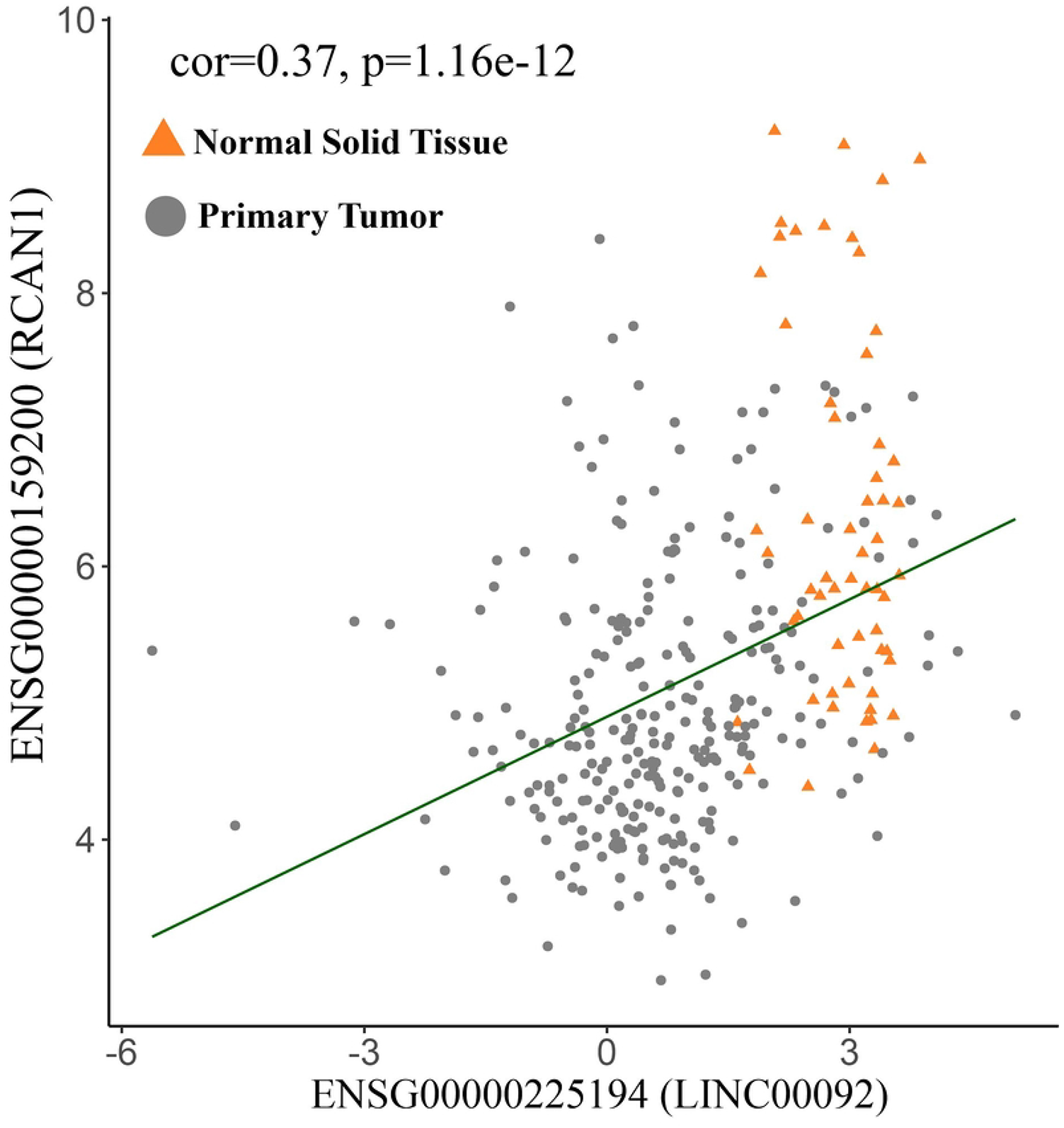
Analysis of the correlation between LINC00092 and RCAN1 based on correlation coefficients.

**Fig 4.**
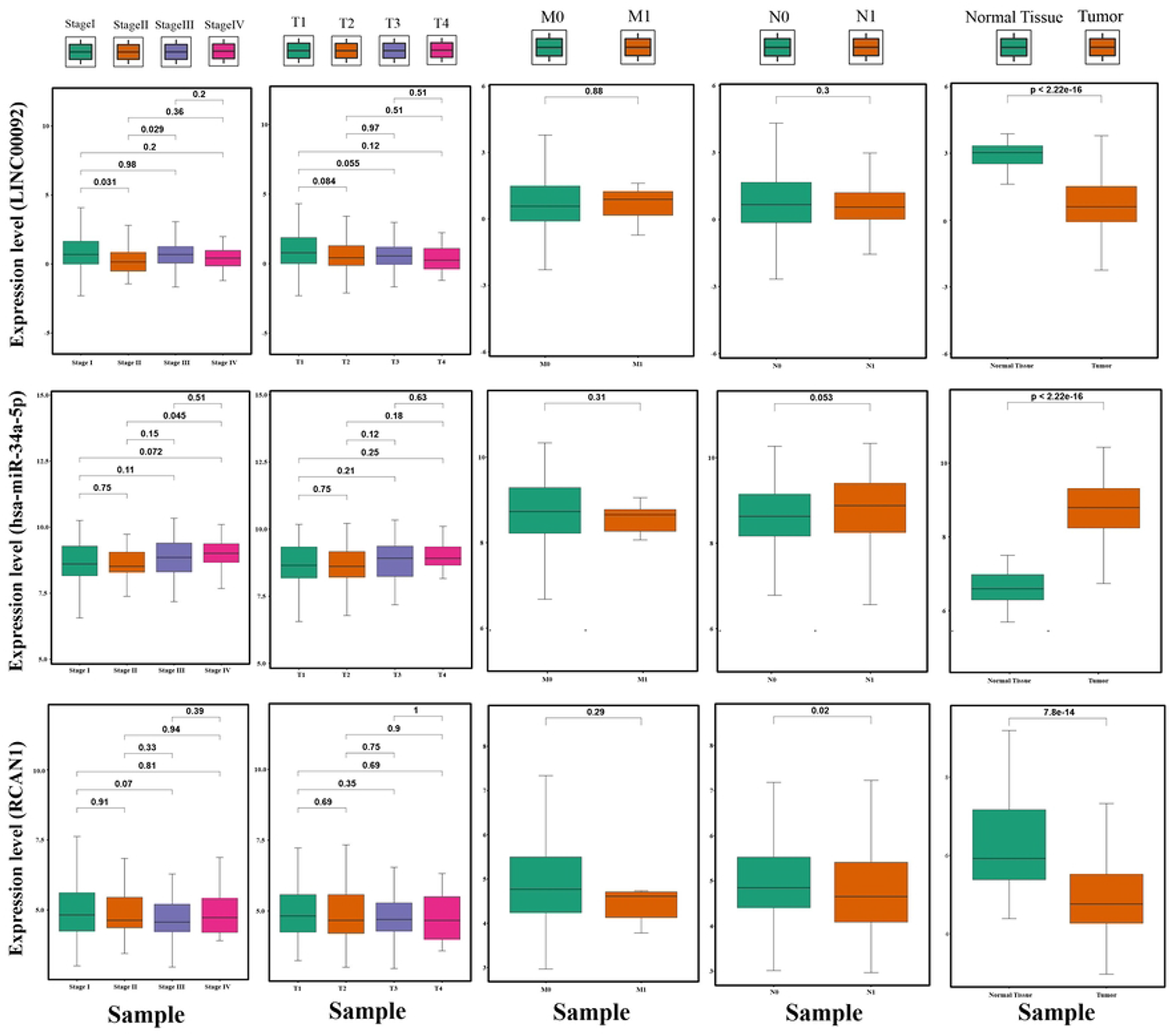
Correlation between expression of LINC00092, has-miR-34a-5p, RCAN1, and TNM stage traits of THCA.

### PPI analysis of RCAN1

Using the GeneMANIA plugin in Cytoscape, Protein-Protein Interactions (PPIs) networks were generated for RCAN1 (Fig 5). In constructing the composite network, each edge (link) in the network is weighted based on individual data sources.

**Fig 5.**
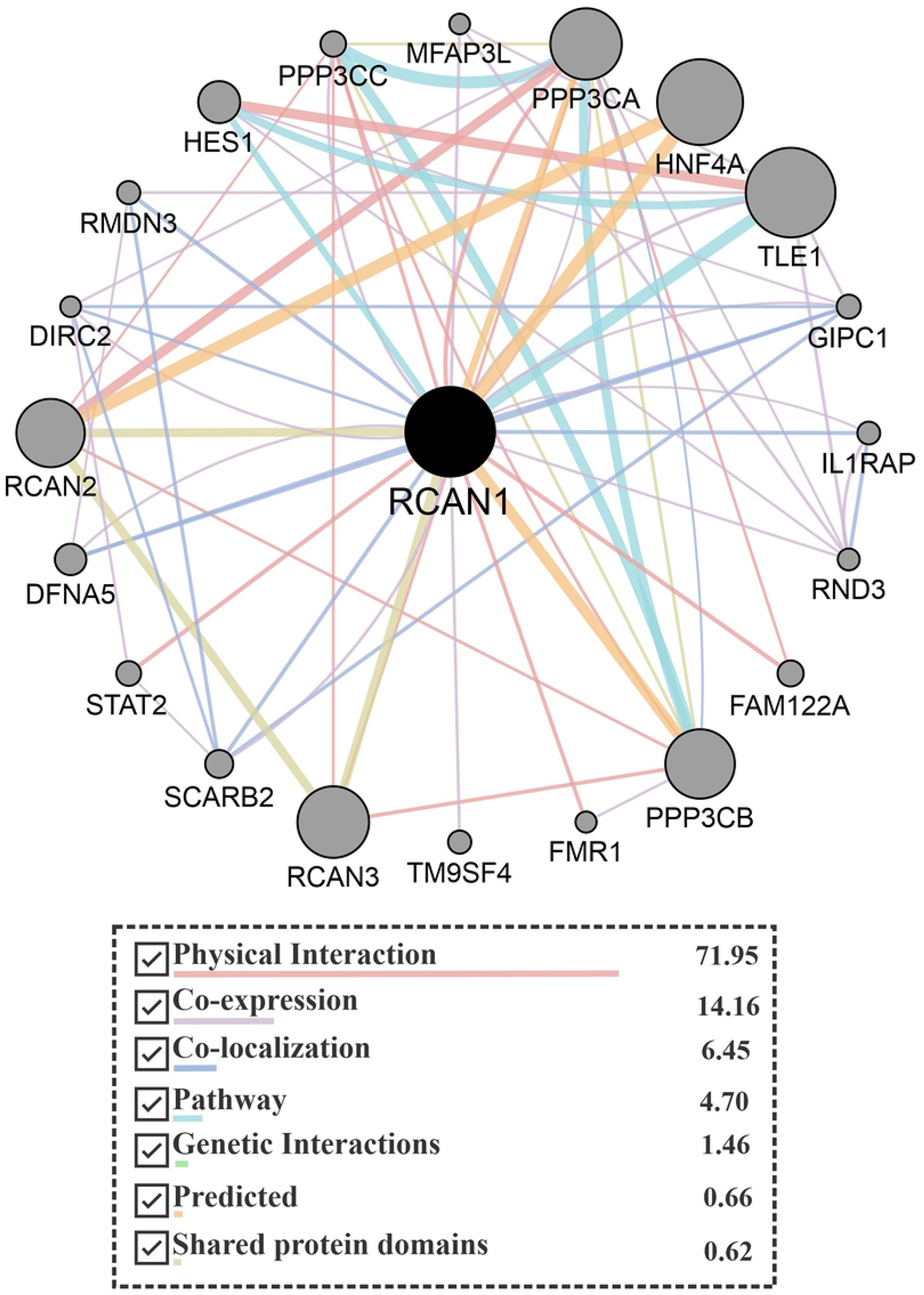
RCAN1 protein-protein interaction network includes 21 nodes and 68 edges. Nodes are sized according to the degree of value of each gene. It is important to note that the colors of the lines represent the type of interaction (Physical interaction, Co-expression, co-localization, pathway, genetic interaction, predicted, shared protein domains). The thickness of a line indicates how strong the support for the data is.

According to the program, the members of this association network were linked through the following networks: physical interactions 71.95 percent, co-expression 14.16 percent, co-localization 6.45 percent, pathways 4.70 percent, genetic interactions 1.46 percent, predicted 0.66%, shared protein domains 0.62%. The network weights add up to 100% and indicate the importance of each data source in predicting query list membership. The genes were ranked using these scores. Each gene pair’s score represents how often pathways that begin at a specific gene node end up in one of the query nodes and how long and intensely weighted those paths.

### Functional Enrichment Analysis of DEGs

The biological functions and signaling mechanisms of genes associated with the RCAN1 PPI network were investigated using Gene Ontology (GO) and Kyoto Encyclopedia of Genes and Genomes (KEGG) enrichment analyses.

GO, and KEGG analyses were performed on RCAN1 PPI network genes. In terms of biological process (BC), the top three GO results indicate the association of genes with “calcium-mediated signaling”, “second-messenger-mediated signaling”, and “calcineurin-NFAT signaling cascade”. In terms of cellular component (CC), the top three GO results indicate genes were enriched in “ glutamatergic synapse”, “magnesium-dependent protein serine/threonine phosphatase complex”, and “protein serine/threonine phosphatase complex”. In terms of molecular function (MF), the top three GO results indicate genes were enriched in “protein phosphatase regulator activity”, “protein serine/threonine phosphatase activity”, and “phosphatase regulator activity”. As a result of KEGG pathway analysis, “Kaposi sarcoma-associated herpesvirus infection”, “C-type lectin receptor signaling pathway”, “Th17 cell differentiation”, “ Osteoclast differentiation”, and “Wnt signaling pathway” emerged as the top five pathways (Fig 6). In order to visualize the enrichment results relating to Biological processes and Molecular functions, the ClusterProfiler package provides a cnetplot function that can be used to plot data (Fig 7).

**Fig 6.**
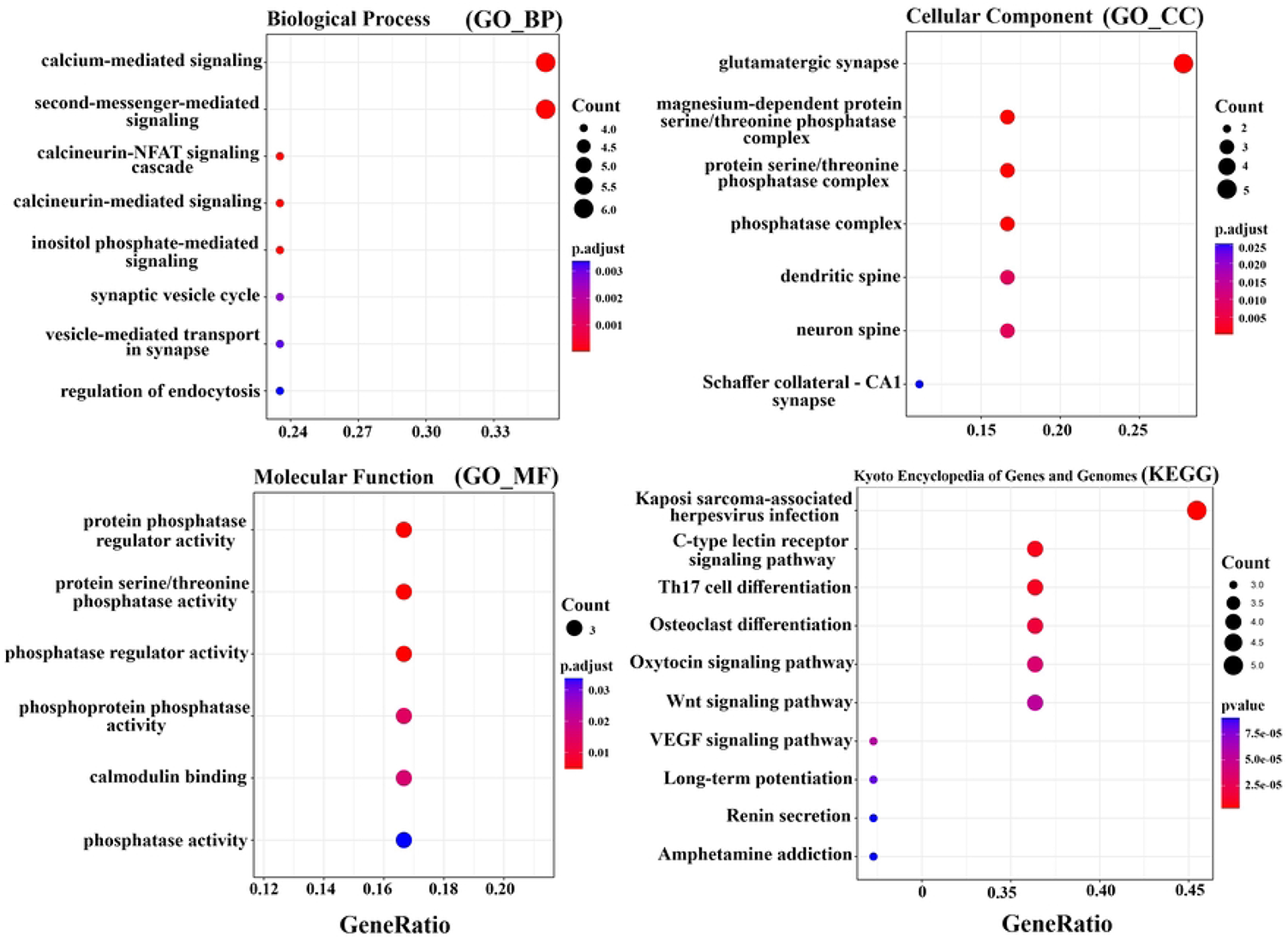
Each dot plot displays an overview of GO and KEGG analysis; each dot plot shows significantly enriched terms in categories of Gene Ontology, including **A)** biologic processes, **B)** cellular components, **C)** molecular functions, and also enriched terms in **D)** KEGG. There is a greater significance to the terms at the top of the dot plot than the terms at the bottom.

**Fig 7.**
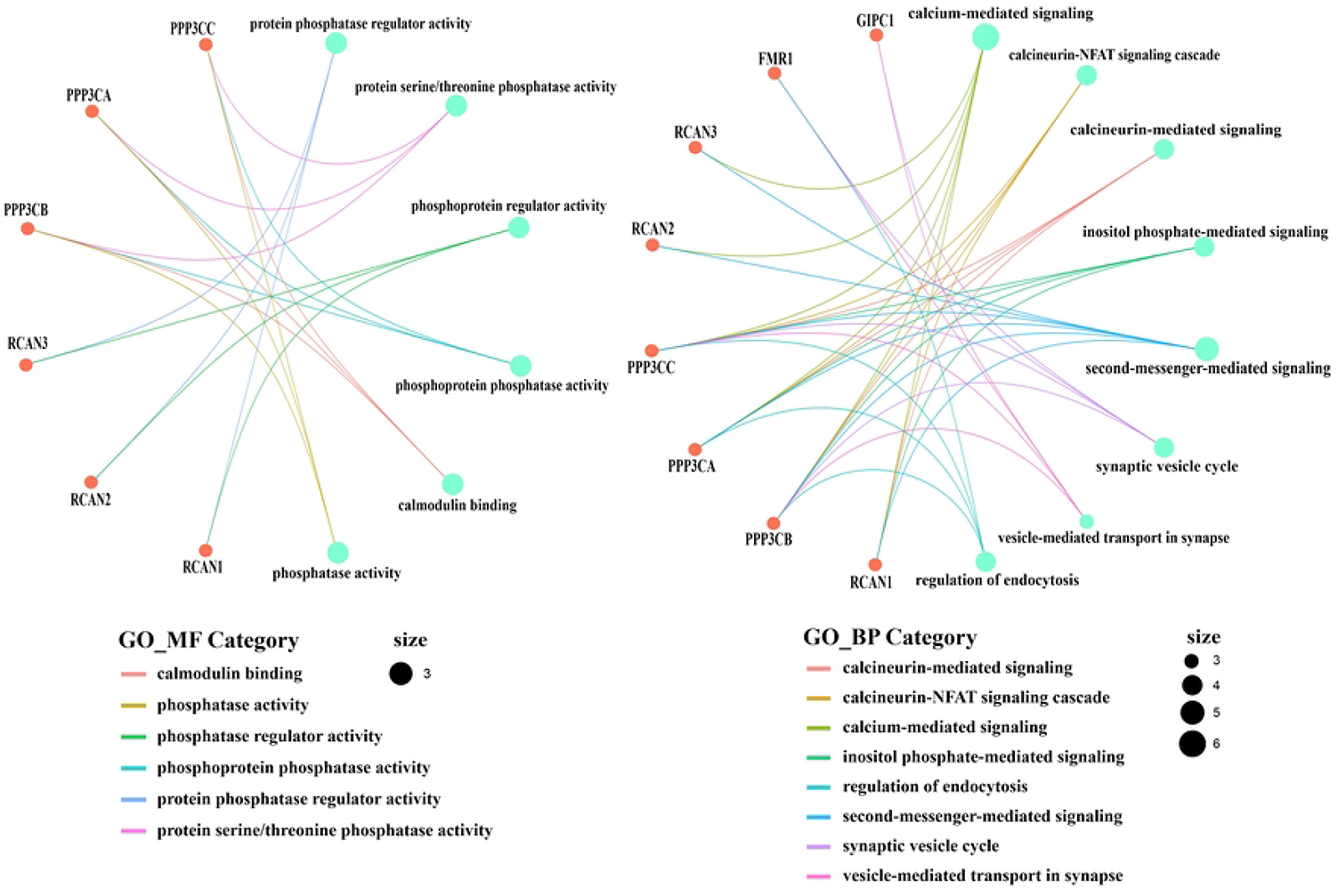
Cnetplot depicts the connection of proteins in the PPI network and GO terms associated with biological process and molecular function.

### RCAN1 Expression is Correlated with Immune Infiltration

One of the cancer hallmarks is the immune reaction. Solid tumors are generally infiltrated with immune cells consisting of B and T lymphocytes, macrophages, neutrophils, dendritic cells (DCs), etc. Infiltration of cells is responsible for chronic inflammation. Increased data demonstrated that local inflammation powerfully induces cancer development. The correlation between RCAN1 and the infiltration levels of immune cells was analyzed by TIMER. In figure 8 the results showed that RCAN1 expression was positively correlated to T cell CD8+ (Rho = 0.285, p = 1.35e−10), Macrophage (Rho = 0.28, p = 3.19e−10) and Neutrophil (Rho = 0.159, p = 4.29e−04) infilteration. However, there was no significant correlation between RCAN1 expression and CD4+ T cell (Rho = −0.066, p = 1.44e−01), B cell (Rho = −0.046, p = 3.11e−01) and Myeloid dendritic cell (Rho = −0.082, p = 6.87e−02) infiltration.

**Fig 8.**
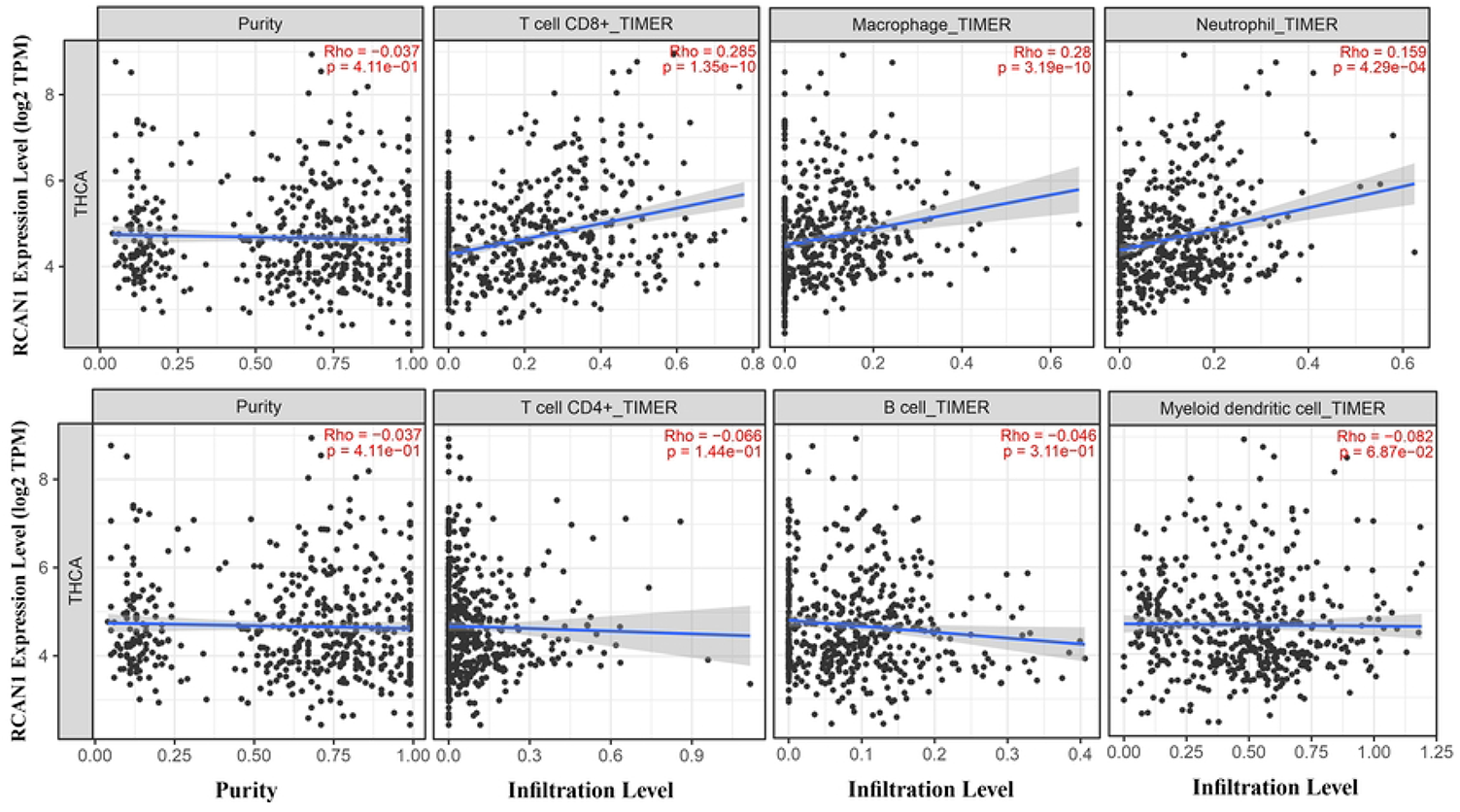
Infiltrations of six immune cells by TIMER are observed to correlate with LINC00092 expression.

## Discussion

Diagnoses and treatments for thyroid cancer patients have evolved over the past several decades based on improved classification of patients, better matching clinical outcomes, and advances in imaging, laboratory, and other techniques (10). In recent years, improvements in thyroid cancer diagnosis and targeted therapies have been made possible by advances in knowledge regarding thyroid cancer’s molecular/genomic characteristics. Based on the expression of the target or a biomarker, there are drugs which are FDA-approved for use in a tumor-agnostic manner (10,40). As a result, a deeper understanding of thyroid cancer’s molecular pathology will further impact thyroid cancer diagnosis and treatment.

In recent years, it has become clear that non-coding RNAs (ncRNA) play an important role in the progression of TC. Nevertheless, the molecular mechanism of this malignancy is unknown mainly due to the complex interactions between multiple kinds of ncRNAs (41). The competitive endogenous RNA (ceRNA) mechanism is a recent discovery that indicates how various RNAs interact and regulate each other. Several ceRNA networks appear to be dysregulated during TC development, metastasis, migration, invasion, epithelial-mesenchymal transition, and drug resistance, based on reviews of relevant literature. Therefore, identifying the dysregulations in TC patients could lead to earlier diagnosis and more effective treatments. According to some studies, some non-coding RNAs (ncRNAs) interact with microRNAs (miRNAs) and modulate gene expression at the transcriptional level and protein translation by sharing microRNA response elements (MRE). In the ceRNA hypothesis, lncRNAs have gained attention for their essential regulatory roles as miRNA decoys (42). Researchers are increasingly proving that lncRNAs play a critical role in the initiation and progression of cancers, including TC. According to recent researches, lncRNA can be detected in plasma, serum, and other fluids with good stability, suggesting new avenues for research on lncRNAs, which can serve as therapeutic targets and prognostic markers (43). Consequently, these studies investigated the significance of lncRNAs as ceRNA correlated with the development and progression of TC.

There is evidence that the downregulation of LINC00092, a long noncoding RNA located at 9q22.32, plays an essential role in cancer development. In lung adenocarcinoma (LUAD), downregulation of LINC00092 contributes directly to tumor progression and metastasis, and depletion of LINC00092 has been associated with poor outcomes (44). Several mechanisms have been identified that might be responsible for LINC00092’s effect on LUAD, including the NF-kB, HIF-1, and ErbB signaling pathways. A study showed that LINC00092 suppressed glycolysis to inhibit human cardiac fibrosis (HCF) activation. This process might be facilitated by LINC00092’s inhibition of extracellular-signaling-regulated kinase (ERK), which justifies why the downregulation of LINC00092 in HCF accelerated proliferation and enhanced migration (45).

The expression of LINC00092 was down-regulated in breast invasive ductal carcinoma (IDC), which was correlated with poor prognosis and promoted cell proliferation, colony formation, cell migration, and invasion. This study found that LINC00092 acts as a sponge for miR-1827. Considering that secreted frizzled-related protein 1 (SFRP1) can directly be targeted by miR-1827, LINC00092/miR-1827/SFRP1 is a signal axis to regulate malignant cell behaviors in IDC (46).

The functional role of LINC00092 in some cancers remains unclear despite the fact that it has been the subject of several studies. The results of our bioinformatics analysis indicate that LINC00092 is capable of suppressing miR-34a-5p. It has been demonstrated that miR-34a-5p has essential biological functions in regulating some tumors; for instance, in Renal Cell Carcinoma (RCC), lncRNA 00312 may stimulate the expression of ASS1 by inhibiting miR-34a-5p’s ability to bind to ASS1 mRNA. Because of this, lncRNA 00312 may exert antitumor activity through normalizing ASS1 expression. ASS1, an enzyme that catalyzes the synthesis of arginine succinic acid by citrulline and aspartic acid, catalyzes the reaction of citrulline with aspartic acid to produce arginine succinic acid, which is targeted by miR-34a-5p and can be negatively regulated by miR-34a-5p. It has been shown that abnormally activated ASS1 contributes to tumor development and tumor growth in RCC (47). In another study, the expression of CASC7, miR-34a-5p, and TP73 were also examined in papillary thyroid cancer (PTC) tissues. There were a significant downregulation of CASC7 and TP73 expression in PTC tissues, while miR-34a-5p expression was elevated. They found that CASC7 activated miR-34a-5p to upregulate TP73 expression, inhibiting proliferation and migration of PTC cells and forcing them to apoptosis (48).

As a result of our analysis, RCAN1 is also one of the targets of miR-34a-5p. It plays a crucial role in cancer pathogenesis as an endogenous inhibitor of calcineurin called Regulator of Calcineurin 1 (RCAN1). RCAN1 downregulation induces cell proliferation, migration, invasion, and angiogenesis while suppressing apoptosis through its loss of inhibitory effect on CN-NFAT (calcineurin-nuclear factor of activated T cells). In several cancers, RCAN1 expression is significantly reduced, including urothelial bladder carcinoma (BLCA), breast invasive carcinoma (BRCA), kidney chromophobe (KICH), kidney renal papillary cell carcinoma (KIRP), liver hepatocellular carcinoma (LIHC), and lung adenocarcinoma (LUAD) (49). According to another study performed on thyroid cancer cells, it was shown that the loss of endogenous RCAN1 elevated the amount of cell proliferation in 3D conditions, increased invasion through Matrigel in vitro, promoted tumor growth in flank xenograft models, and increased tumor metastasis (50).

We have found that the downregulation of LINC00092 plays a functional role in the progression of TC based on the results of our analysis. Further analysis was also conducted on the expression patterns of LINC00092 in 31 different cancer types. Figure 9 indicates that LINC00092 was significantly downregulated than normal tissues in Cervical squamous cell carcinoma and endocervical adenocarcinoma (CESC), Brain Lower Grade Glioma (LGG), Lung adenocarcinoma (LUAD), Lung squamous cell carcinoma (LUSC), Ovarian serous cystadenocarcinoma (OV), Stomach adenocarcinoma (STAD), Testicular Germ Cell Tumors(TGCT) and Thyroid carcinoma (THCA).

**Fig 9.**
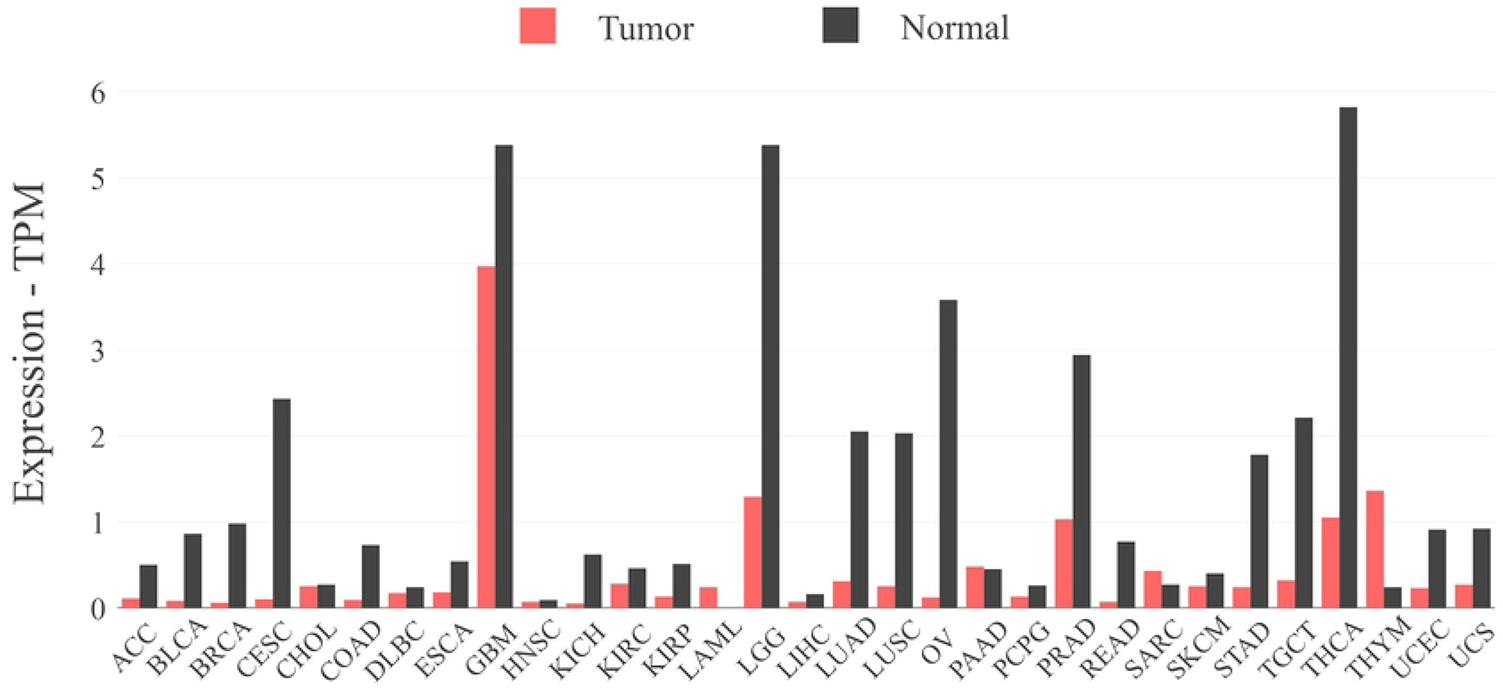
LINC00092 expression patterns are observed in 31 types of cancer.

According to our study results, LIN00092 inhibits miR-34a-5p expression in normal thyroid tissue, which enables RCAN1 to continue to function as a tumor suppressor. On the other hand, when the expression of LINC00092 is reduced in thyroid cancer tissue, miR-34a-5p is increased. As a result, the high level of miR-34a-5p in cancer cells inhibits RCAN1 and decreases its expression, promoting TC proliferation and invasion and inhibiting apoptosis (Fig 10). Our study demonstrated that the LINC00092/miR-34a-5p/RCAN1 axis might play a functional role in the progression of TC. Dysregulation of LINC00092 is a prognostic factor candidate for TC, which provides new insights for TC clinical treatments.

**Fig 10.**
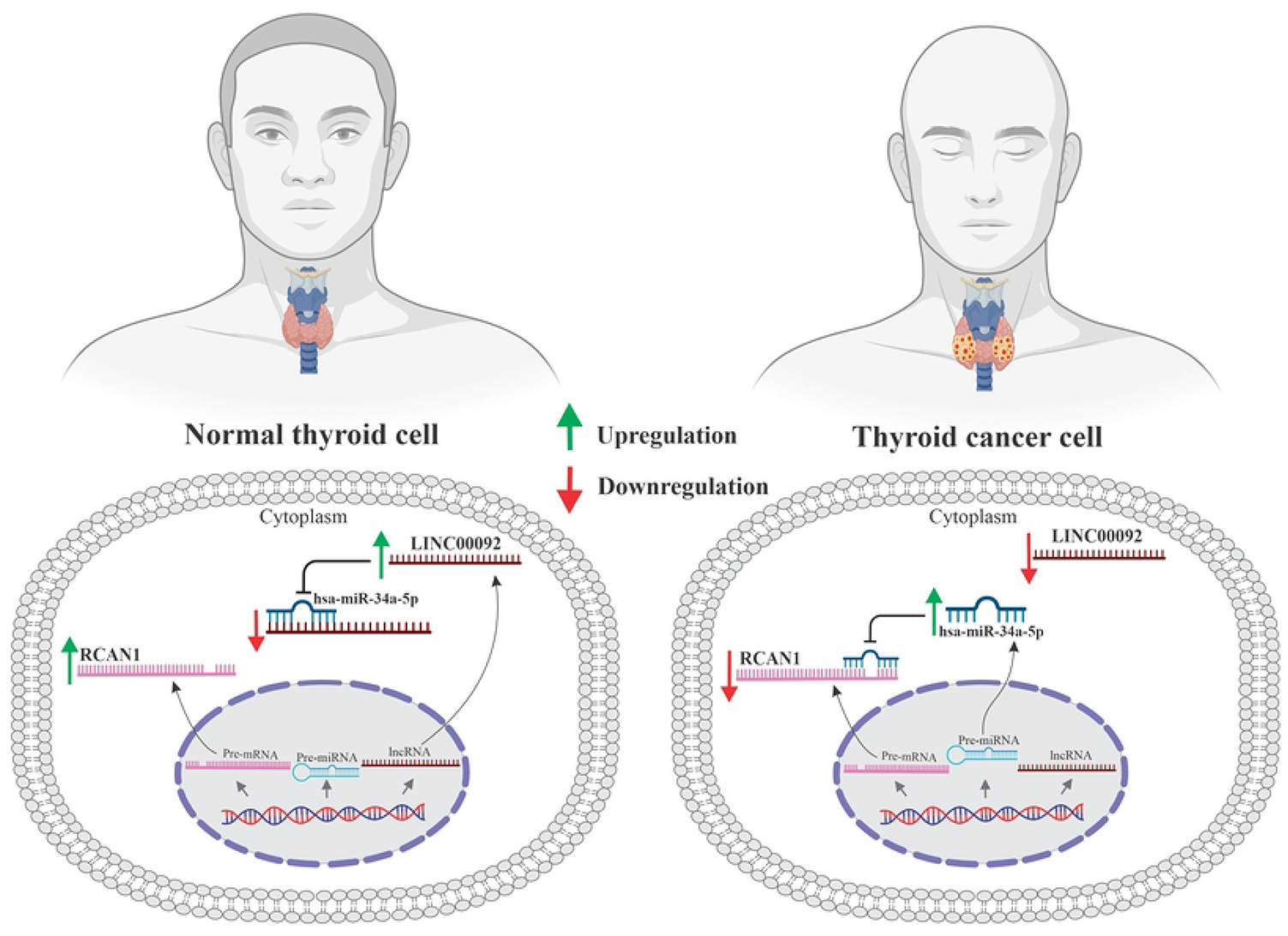
According to our results, LIN00092 inhibits miR-34a-5p expression in normal thyroid cells, thereby maintaining the tumor suppressor function of RCAN1. In contrast, miR-34a-5p levels increase in thyroid cancer cells when LINC00092 expression reduces. Thus, miR-34a-5p inhibits RCAN1 expression in cancer cells, inhibiting apoptosis and promoting TC proliferation and invasion.

## Conclusion

MiR-34a-5p and RCAN1 have been investigated and shown to play a role in thyroid cancer in previous studies. This study aimed to introduce lncRNA LINC00092 as a regulatory molecule capable of modulating the miR-34a-5p/RCAN1 axis. In summary, this study demonstrates that LINC00092 functions as a ceRNA for miR-34a-5p by upregulating its target RCAN1, thus inhibiting cancer progression. According to our results, the LINC00092/miR-34a-5p/RCAN1 axis may play an important role in the progression of TC. Our study aims to provide a deeper understanding of thyroid cancer’s molecular pathology and help thyroid cancer diagnosis and treatment in the future.

## Acknowledgments

We wish to acknowledge The Cancer Genome Atlas (TCGA) project, miRTarBase, DIANA, GEPIA2, Cytoscape, GO, KEGG, and GeneMANIA databases, and their contributors for presenting these valuable public data sets.

## Authors contribution

**SM and PM** wrote the manuscript comprehensively in all parts, **MJK** accompanied in many other sections of the paper and **SHT** critically reviewed the manuscript, edited the manuscript scientifically and technically. All the authors read the manuscript comprehensively and confirmed the final edited version.

